## Supplementary Material for "TT-Mars: Structural Variants Assessment Based on Haplotype-resolved Assemblies"

#### 1 Assembly quality control

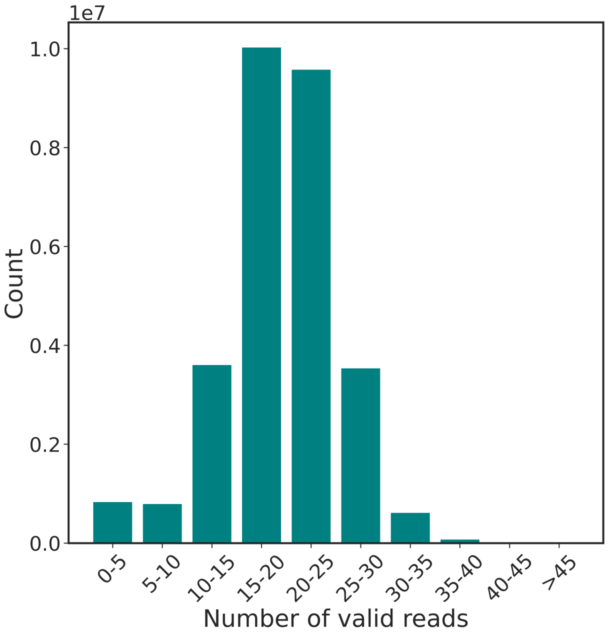


**(a)** Assembly score haplotype 1

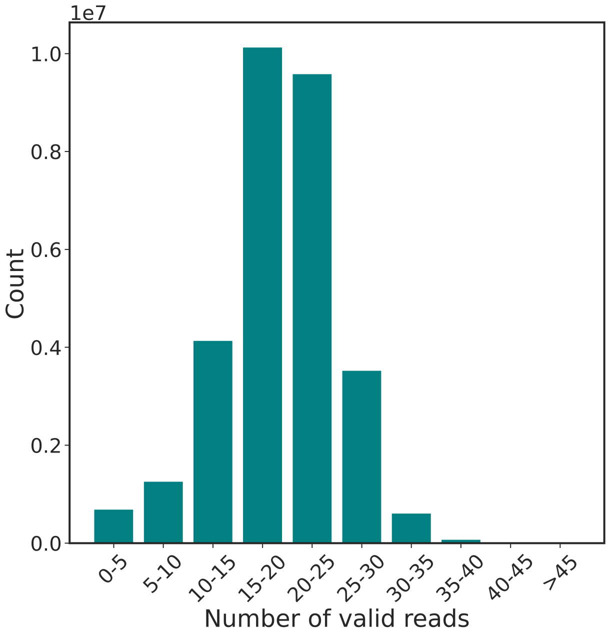


**(b)** Assembly score haplotype 2

**Figure 1:** The distribution of assembly scores across the sample genome HG002, defined by contiguity of sequence alignment. Reads are mapped back to the assemblies. The genome is divided into 100-base bins. A read that spans a bin with at least 1kb flanking length on both sides is considered as a valid read. We count the number of valid reads as the quality score for each bin. Noisy alignment regions and low-coverage regions are given low quality scores.

#### 2 SV operation

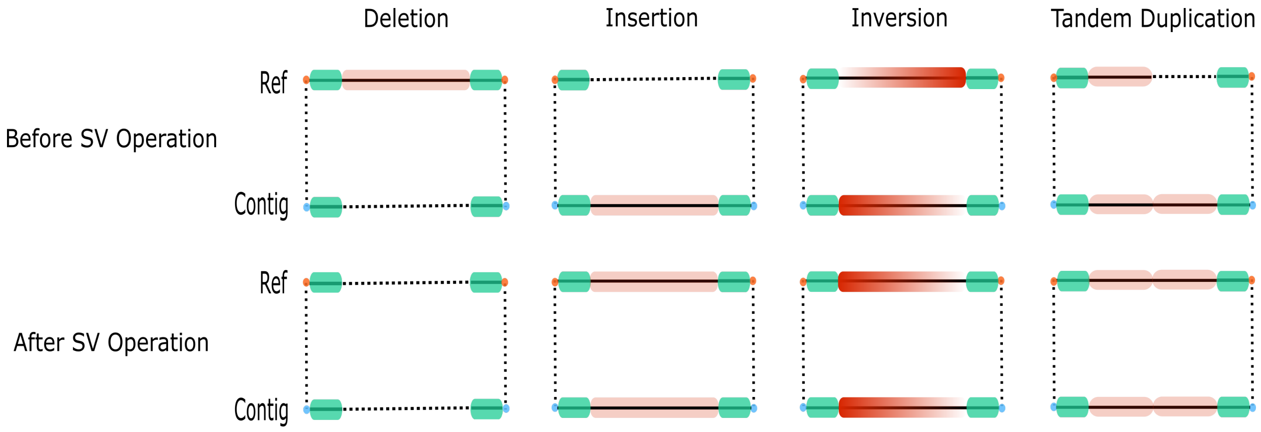


**Figure 2:** An illustration of the classes of SV operations considered by TT-Mars, and the result of the reference before and after each operation. Because the reference is not modified to validate an interspersed duplication, it is not shown here.

#### 3 Dipcall+truvari and TT-Mars classification results loci

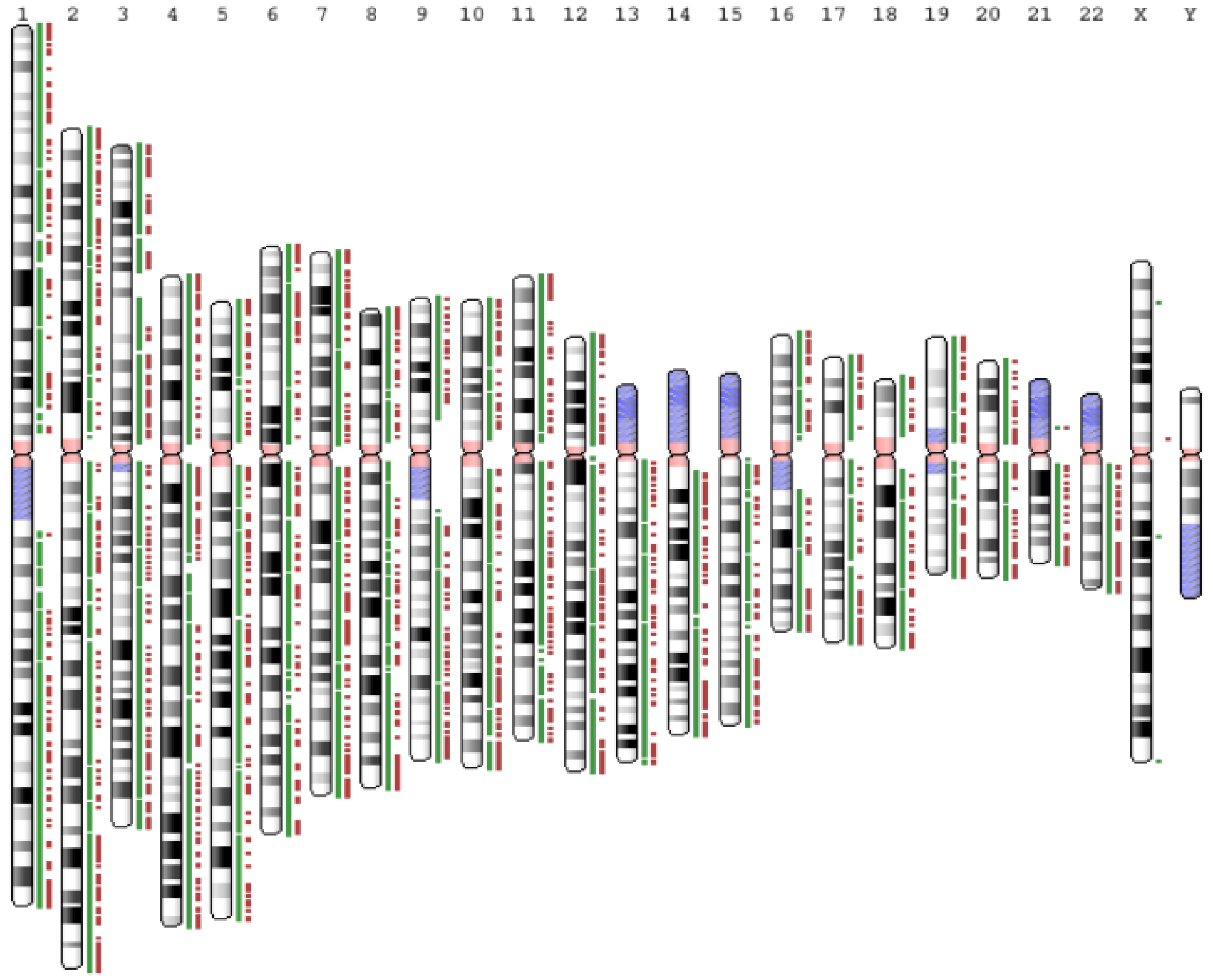


**Figure 3:** Loci of dipcall+truvari and TT-Mars calls from the HG00096 pbsv call set. Green bars are the calls that two method agree on, red bars are the calls that two method disagree on.

### 4 Curation of dipcall+truvari and TT-Mars SV annotation

On the pbsv call set of sample HG00096, we randomly selected 100 calls that are given different results by dipcall+truvari and TT-Mars (50 calls dipcall+truvari FP TT-Mars TP and 50 calls dipcall+truvari TP TT-Mars FP) for manual inspection in IGV. In each figure, the first track is the pbsv call set, the second track is the dipcall call set. The following two tracks are alignments used to generate the orthology map in TT-Mars (lra), the next two tracks are alignments used by dipcall (pbmm2). The track at bottom is the tandem duplication regions.

#### 4.1 Dipcall+Truvari FP and TT-Mars TP results

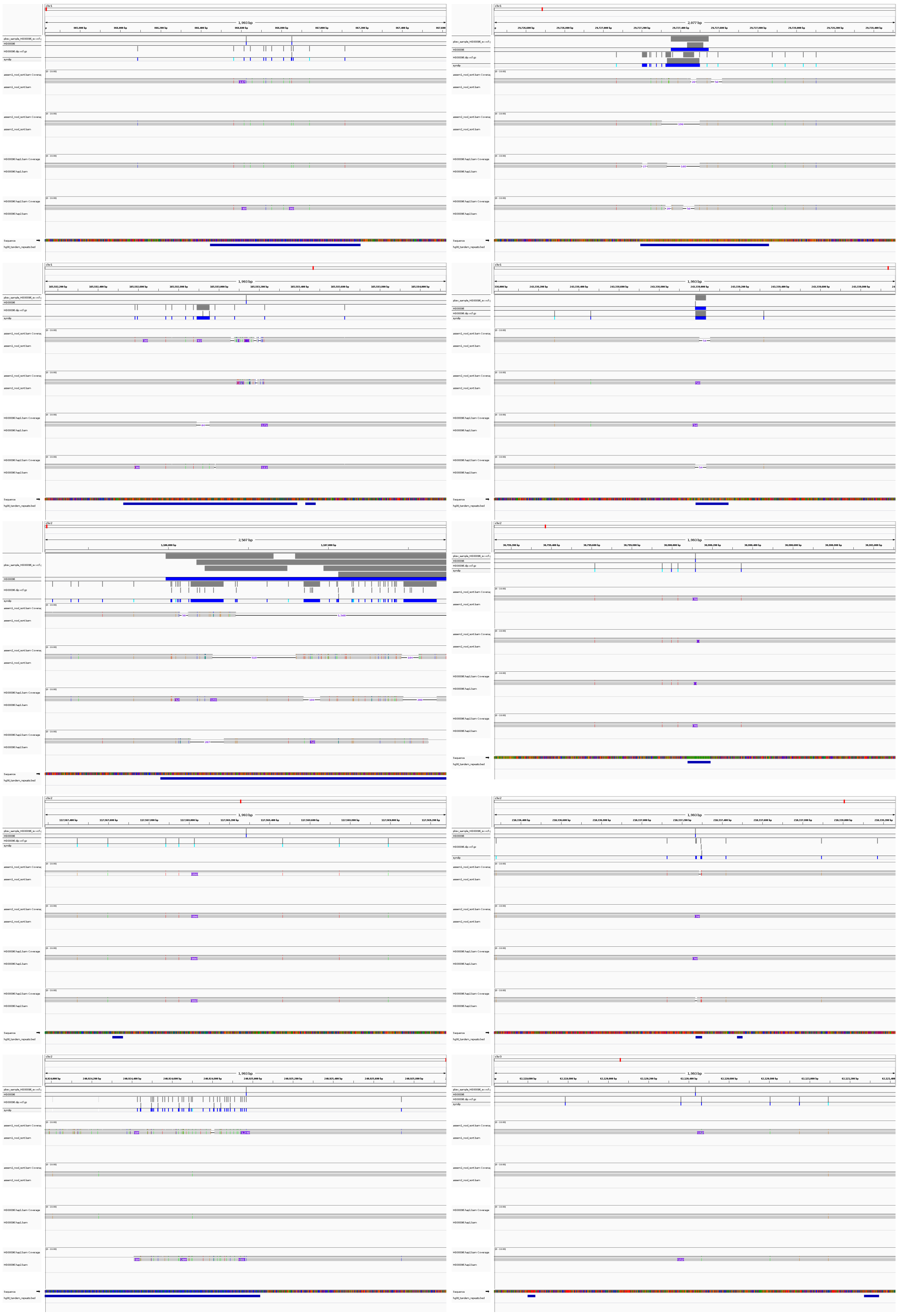

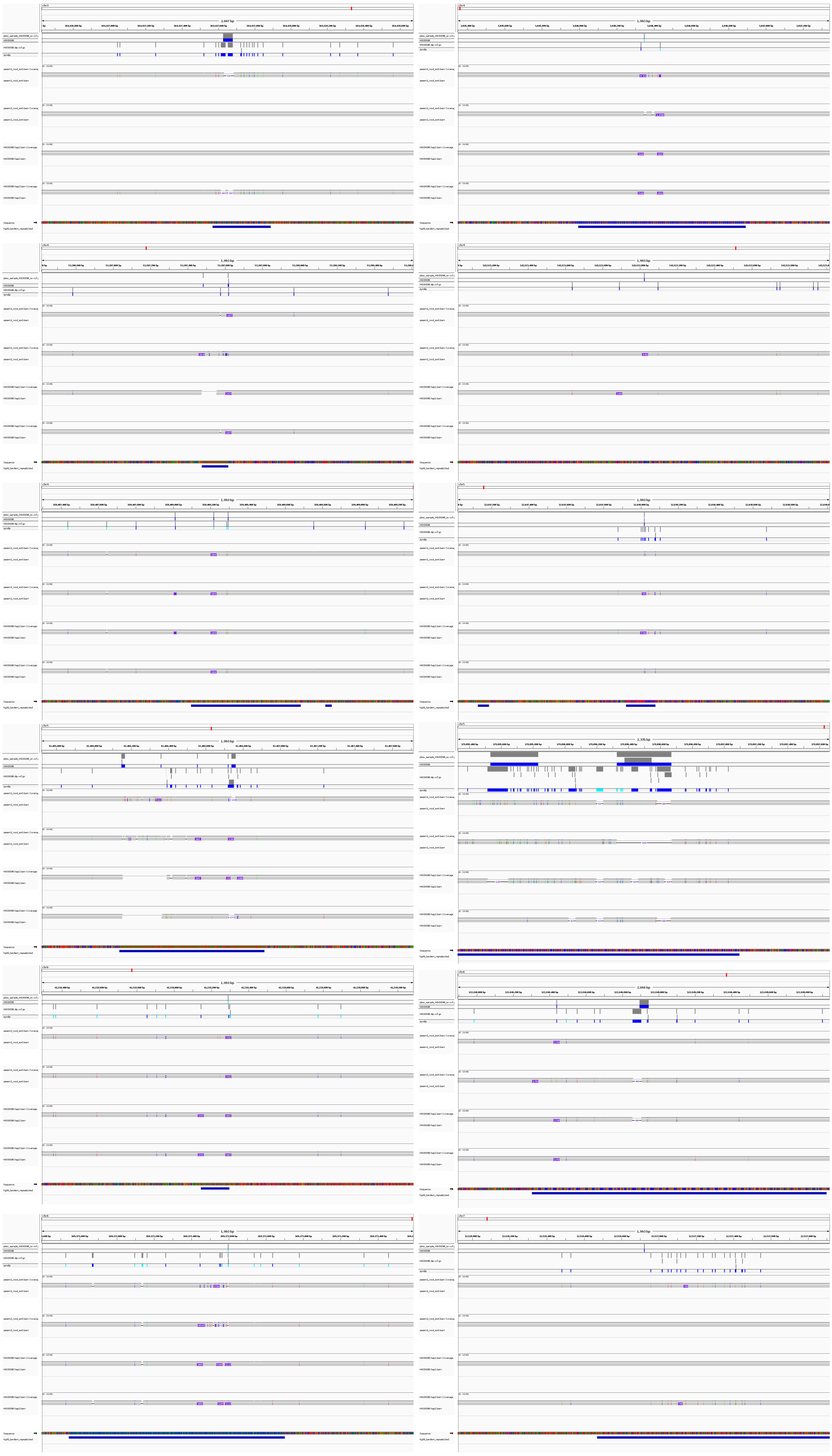

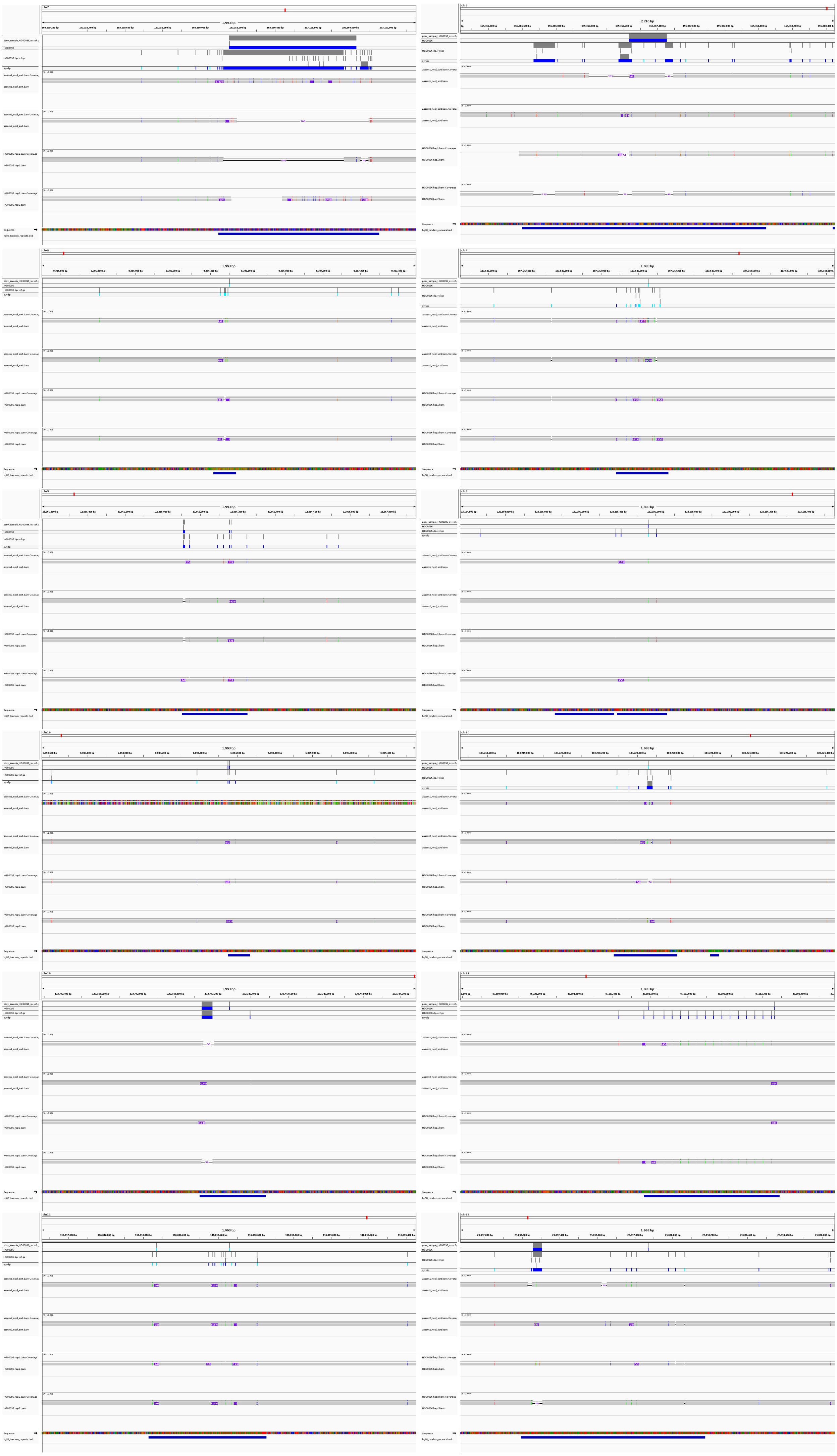

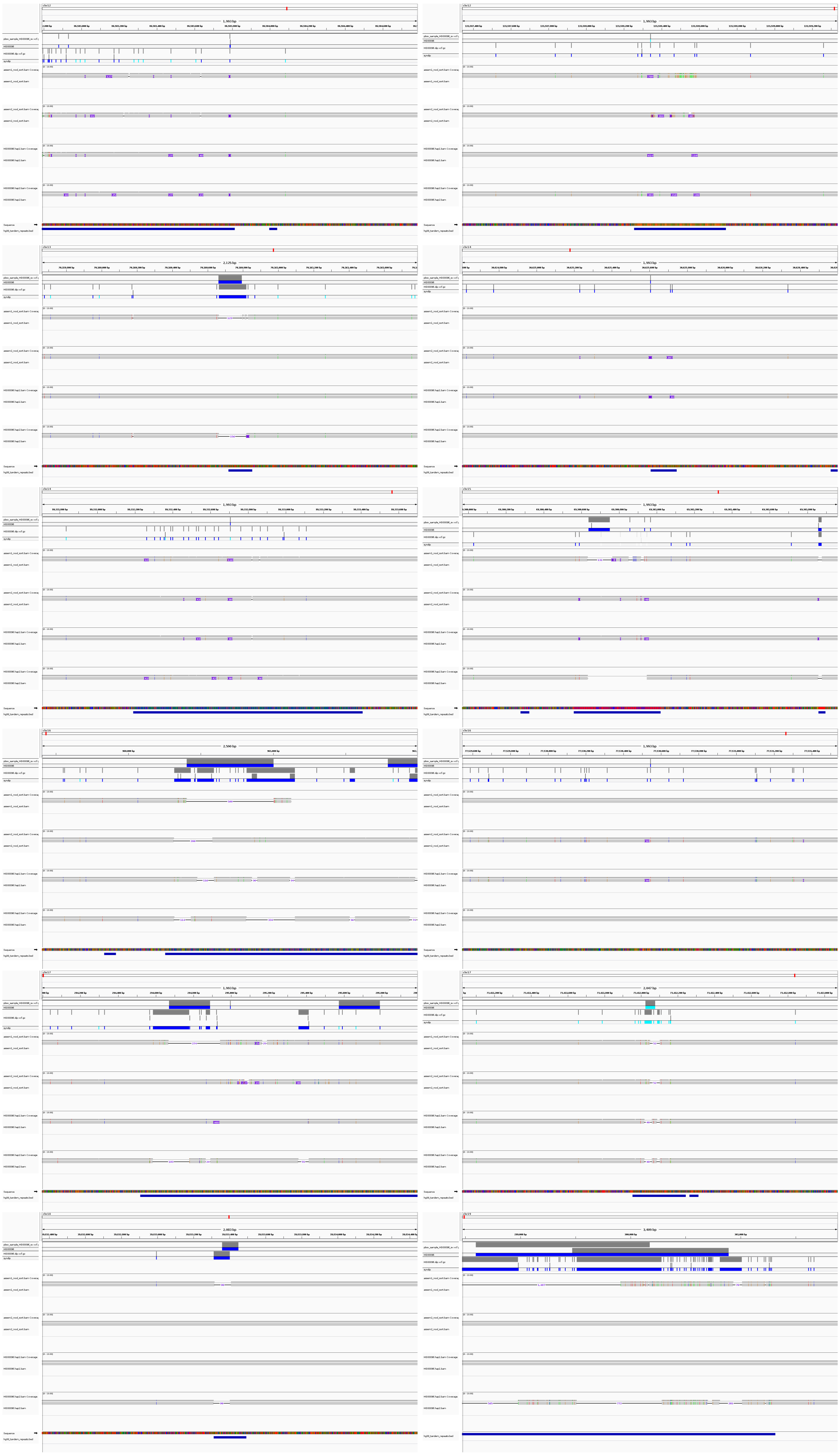

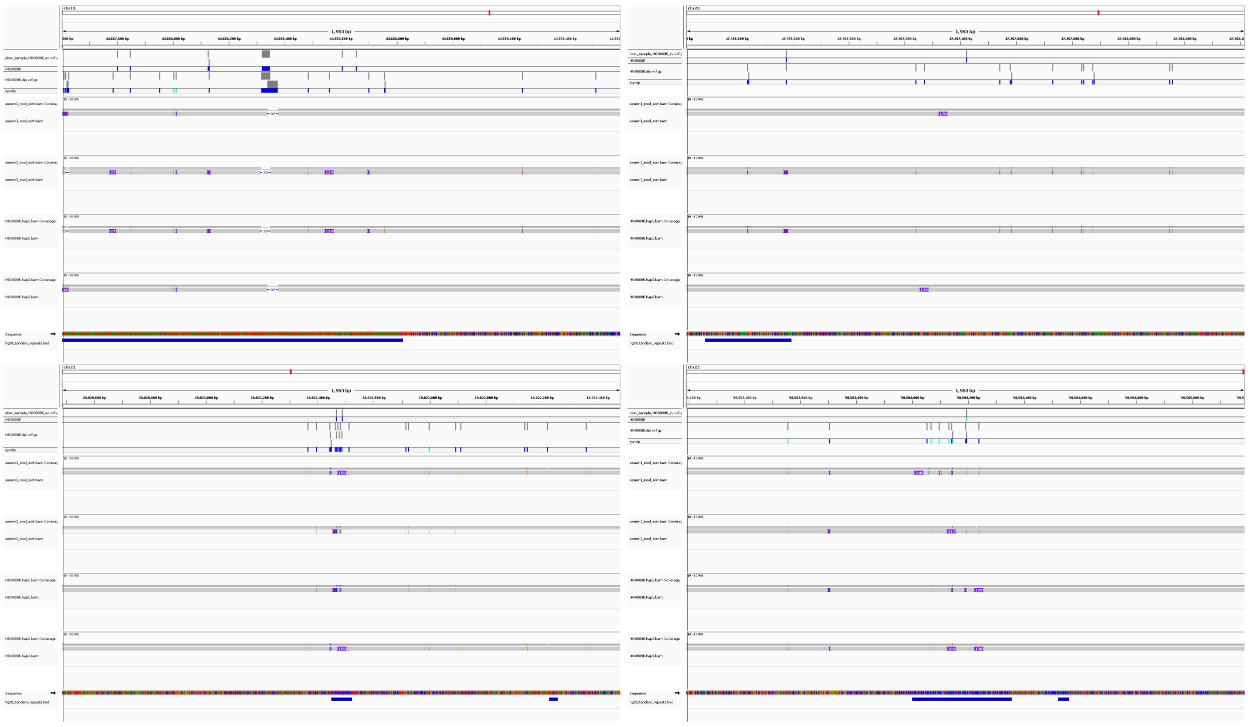

#### 4.2 Dipcall+Truvari TP and TT-Mars FP results

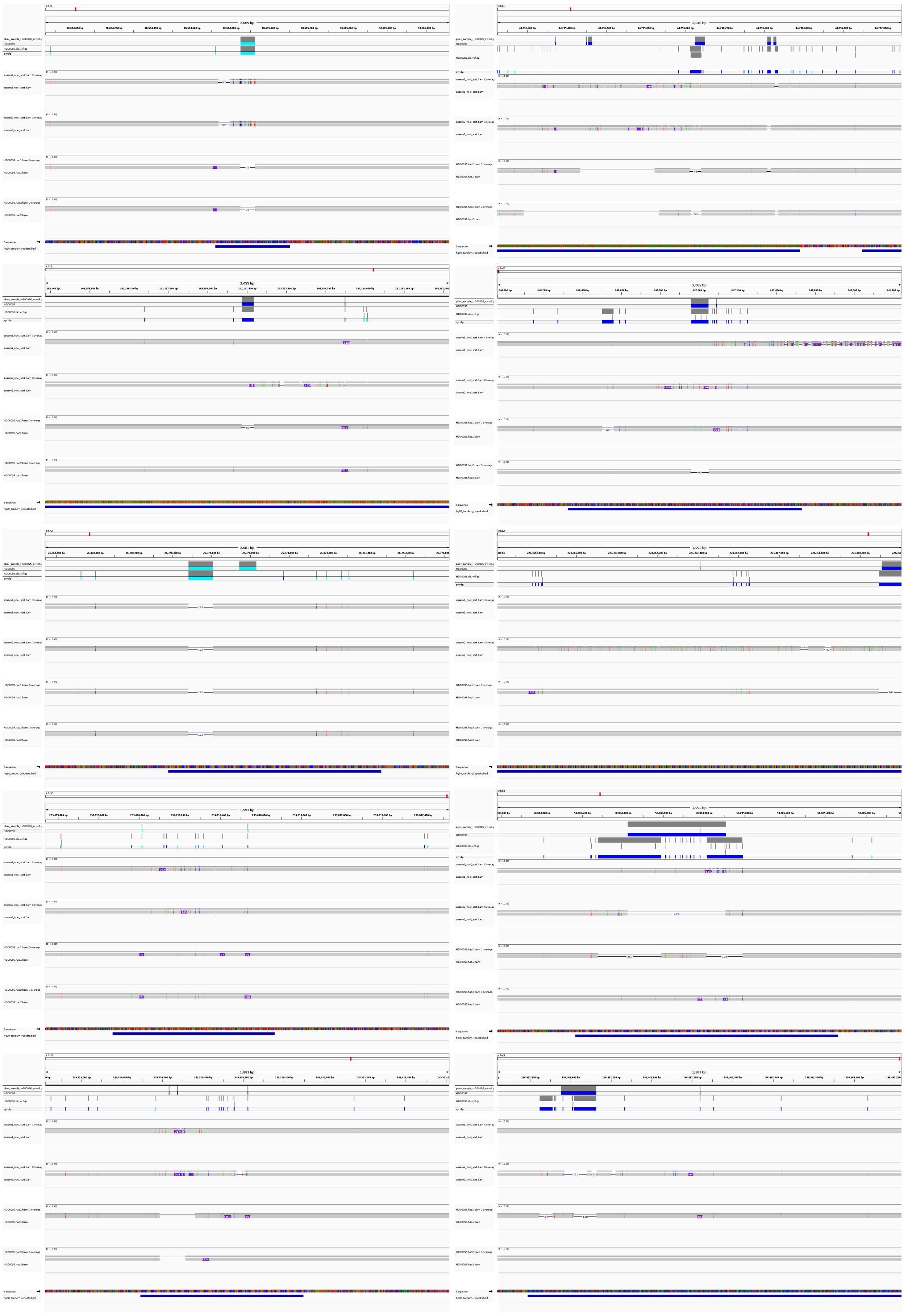

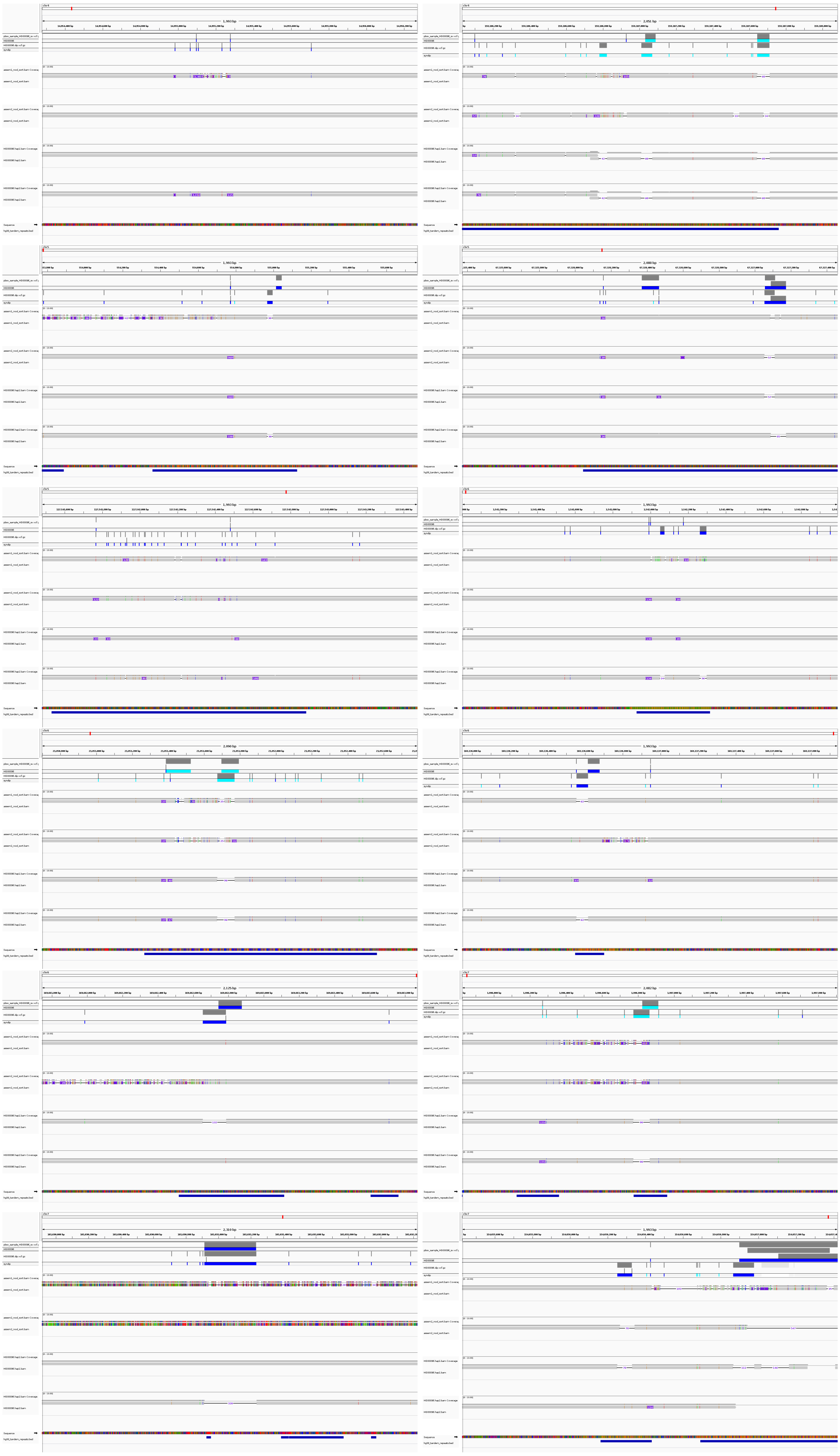

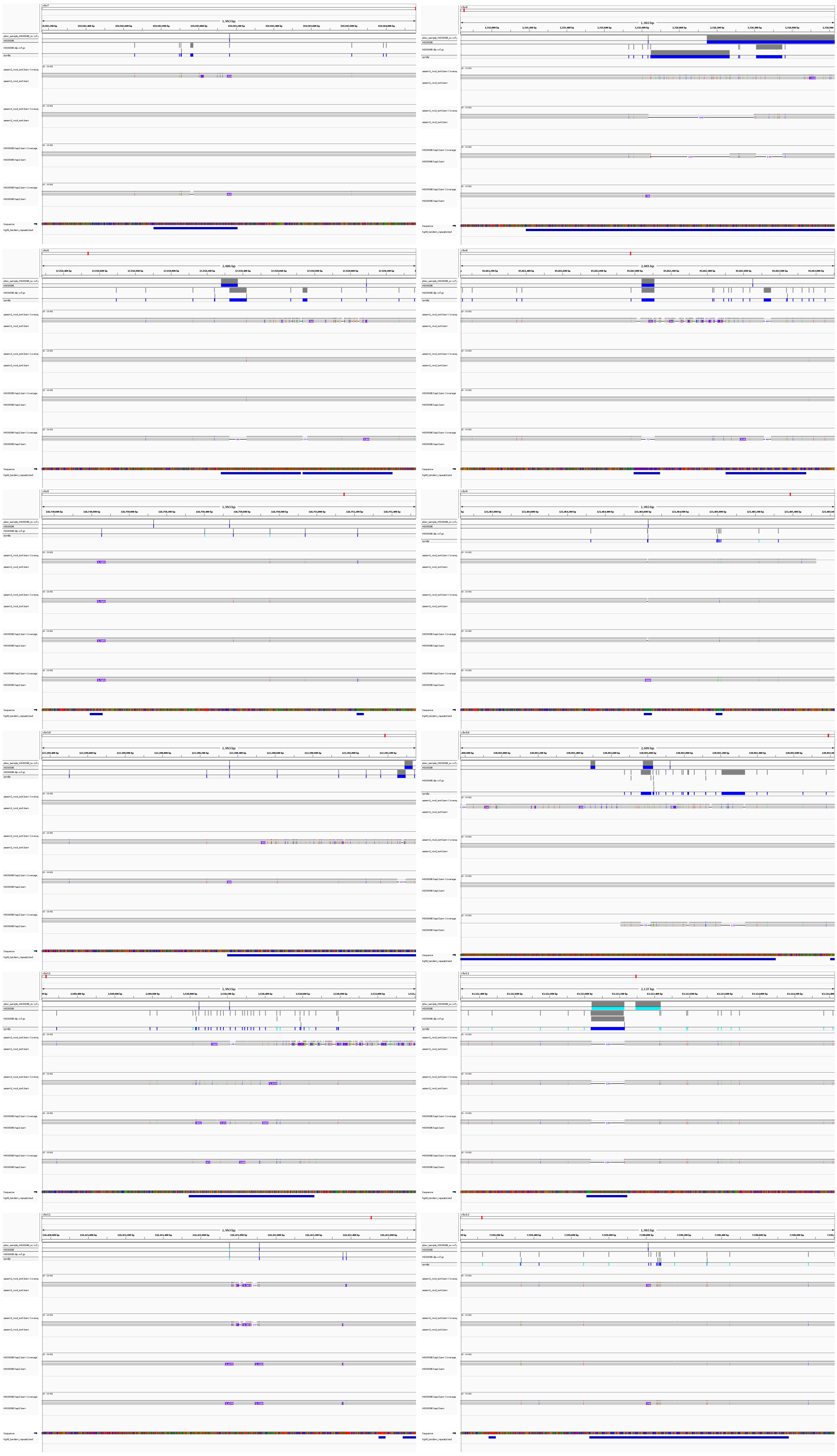

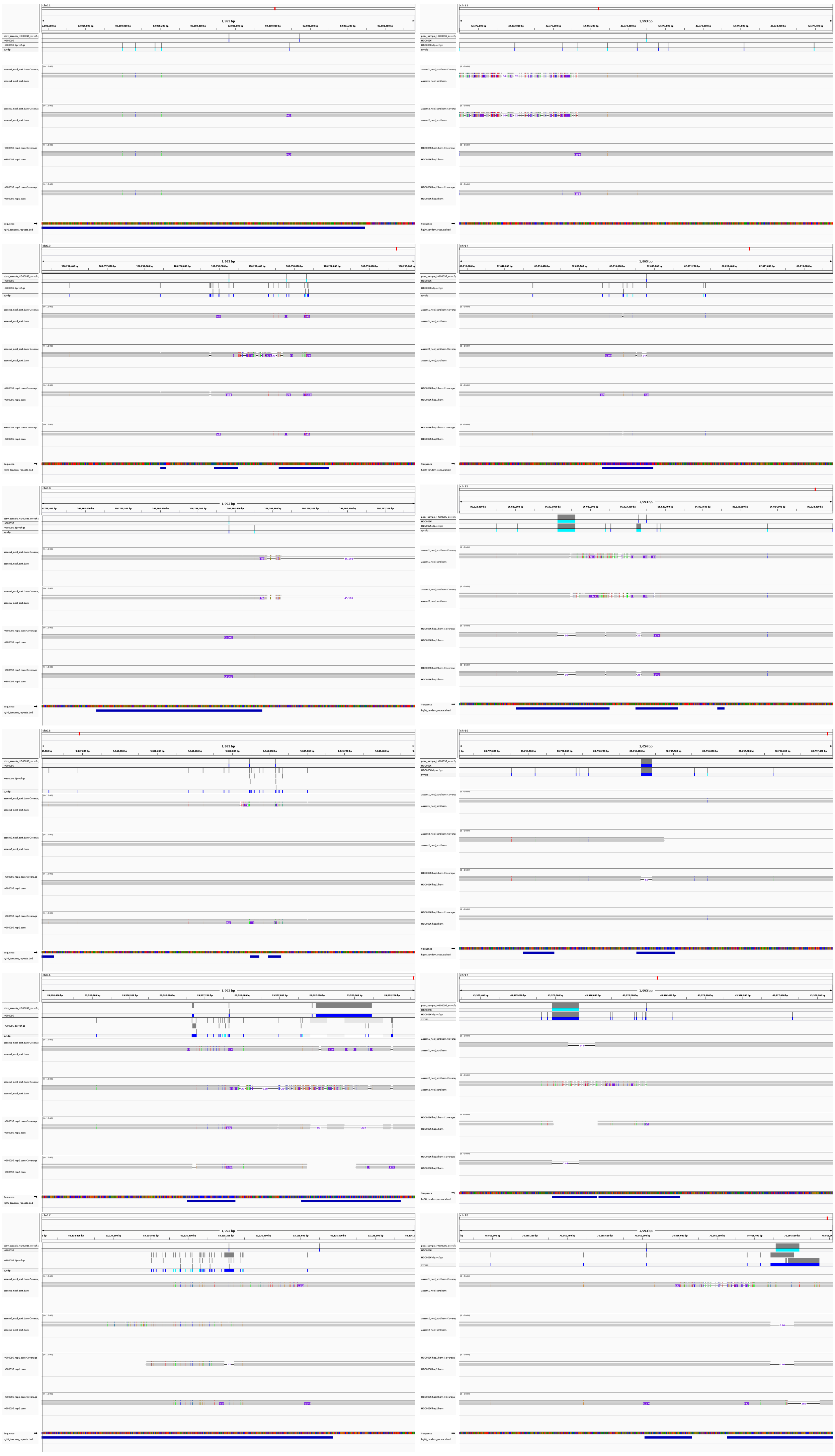

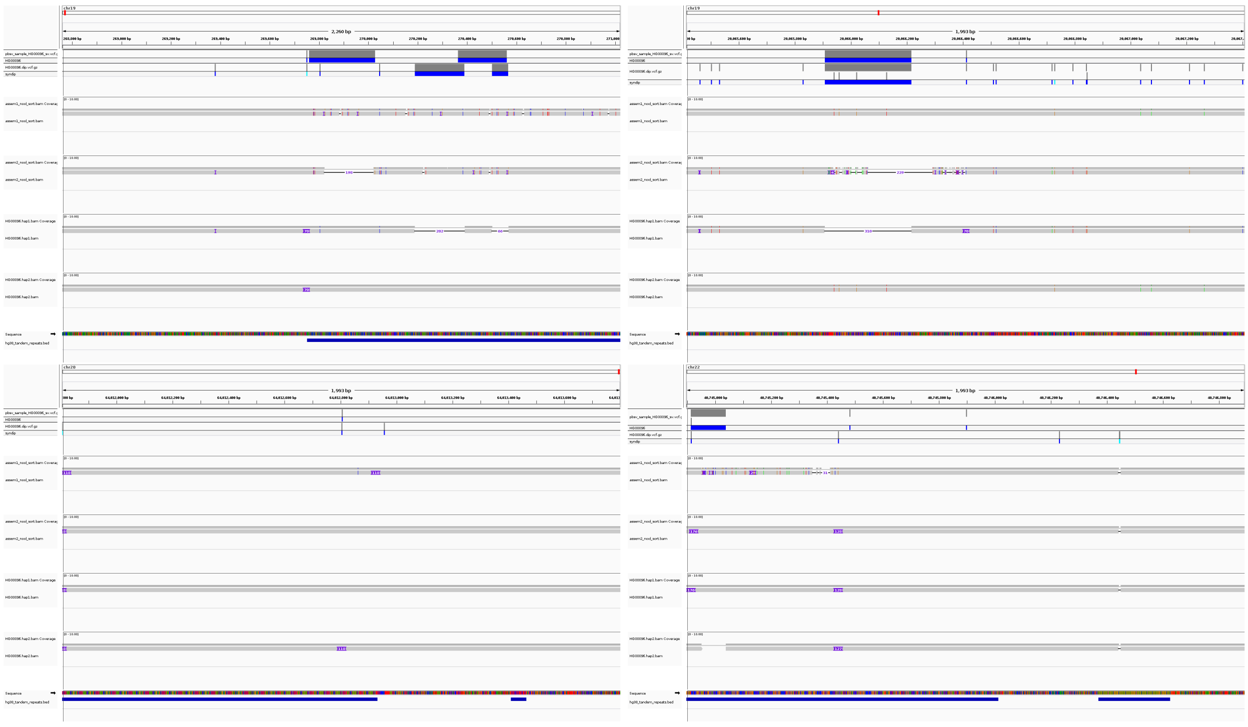

#### 5 Command used to run the callers, TT-Mars and truvari

##### 5.1 Callers

Wham: we followed the suggested command.

```
export EXCLUDE=GL000207.1,GL000226.1,GL000229.1,GL000231.1,GL000210.1,\
GL000239.1,GL000235.1,GL000201.1,GL000247.1,GL000245.1,GL000197.1,\
GL000203.1,GL000246.1,GL000249.1,GL000196.1,GL000248.1,GL000244.1,\
GL000238.1,GL000202.1,GL000234.1,GL000232.1,GL000206.1,GL000240.1,\
GL000236.1,GL000241.1,GL000243.1,GL000242.1,GL000230.1,GL000237.1,\
GL000233.1,GL000204.1,GL000198.1,GL000208.1,GL000191.1,GL000227.1,\
GL000228.1,GL000214.1,GL000221.1,GL000209.1,GL000218.1,GL000220.1,\
GL000213.1,GL000211.1,GL000199.1,GL000217.1,GL000216.1,GL000215.1,\
GL000205.1,GL000219.1,GL000224.1,GL000223.1,GL000195.1,GL000212.1,\
GL000222.1,GL000200.1,GL000193.1,GL000194.1,GL000225.1,GL000192.1,NC_007605
whamg -e $EXCLUDE -a ref.fa -f sample.bam | perl utils/filtWhamG.pl > sample.vcf
2> sample.err
```

LUMPY: we followed the suggested command.

```
samtools view -b -F 1294 sample.bam > sample.discordants.unsorted.bam
samtools view -h sample.bam\
| scripts/extractSplitReads_BwaMem -i stdin\
| samtools view -Sb - > sample.splitters.unsorted.bam
samtools sort sample.discordants.unsorted.bam sample.discordants
samtools sort sample.splitters.unsorted.bam sample.splitters
lumpyexpress -B sample.bam -S sample.splitters.bam\
-D sample.discordants.bam -o sample.vcf
```

DELLY: we followed the default command.

```
delly call -x human.hg38.excl.tsv -o sample.bcf -g ref.fa sample.bam
```

pbsv: we followed the default command for sample HG002.

```
pbsv call ref/human_hs37d5.fasta tools/pbsv/*.svsig.gz tools/pbsv/hg2.pbsv.vcf --ccs
bgzip tools/pbsv/hg2.pbsv.vcf
tabix tools/pbsv/hg2.pbsv.vcf.gz
```

dipcall: we followed the default command.

For female samples:

```
dipcall.kit/run-dipcall prefix ref.fa paternal.fa.gz maternal.fa.gz > prefix.mak
make -j2 -f prefix.mak
```

For male samples, PARs on the reference chrY are hard masked:

```
dipcall.kit/run-dipcall -x dipcall.kit/hs38.PAR.bed prefix ref.fa paternal.fa.gz\
maternal.fa.gz > prefix.mak
make -j2 -f prefix.mak
```

##### 5.2 TT-Mars

TT-Mars:

```
sample=HG00096
reference=path-to-reference_file/hg38.no_alts.fasta
vcf_file=path-to-target-vcf/callset.vcf
asm_h1=path-to-assemblies/h1.fa
asm_h2=path-to-assemblies/h2.fa
output_dir=output-directory
files_dir=./ttmars_files/$sample
centro_file=centromere_hg38.txt
tr_file=hg38_tandem_repeats.bed
if_hg38=True
pass_only=True
seq_resolved=False
num_X_chr=1

python ttmars.py $output_dir $if_hg38 $centro_file\
$files_dir/assem1_non_cov_regions.bed $files_dir/assem2_non_cov_regions.bed \
$vcf_file $reference $asm_h1 $asm_h2\
$files_dir/lo_pos_assem1_result_compressed.bed\
$files_dir/lo_pos_assem2_result_compressed.bed\
$tr_file $pass_only $seq_resolved

python reg_dup.py $output_dir $if_hg38 $centro_file\
```

```
$files_dir/assem1_non_cov_regions.bed $files_dir/assem2_non_cov_regions.bed\
$vcf_file $reference $asm_h1 $asm_h2\
$files_dir/lo_pos_assem1_result_compressed.bed\
$files_dir/lo_pos_assem2_result_compressed.bed $tr_file\
$files_dir/lo_pos_assem1_0_result_compressed.bed\
$files_dir/lo_pos_assem2_0_result_compressed.bed $pass_only

    python chrx.py $output_dir $if_hg38 $centro_file\
$files_dir/assem1_non_cov_regions.bed $files_dir/assem2_non_cov_regions.bed\
$vcf_file $reference $asm_h1 $asm_h2\
$files_dir/lo_pos_assem1_result_compressed.bed\
$files_dir/lo_pos_assem2_result_compressed.bed\
$tr_file $pass_only $seq_resolved

    python combine.py $output_dir $num_X_chr
```

##### 5.3 Truvari

Truvari:

For short-read callers:

```
truvari bench -f /reference/human_hs37d5.fasta --includebed\
HG002_SVs_Tier1_v0.6.bed -b HG002_SVs_Tier1_v0.6.vcf.gz -c HG002.vcf.gz -o\
./HG002/ --sizemax 10000000 --passonly --pctsim=0 -r 1000 --multimatch &
```

For long-read callers:

```
truvari bench -f /reference/human_hs37d5.fasta --includebed\
HG002_SVs_Tier1_v0.6.bed -b HG002_SVs_Tier1_v0.6.vcf.gz -c HG002.vcf.gz -o\
./HG002/ --sizemax 10000000 --passonly -r 1000 --multimatch &
```
